# Complex genetic network underlying the convergent of Rett Syndrome like (RTT-L) phenotype in neurodevelopmental disorders

**DOI:** 10.1101/2020.01.11.899658

**Authors:** Eric Frankel, Julius Dodson, Megan Sharifi, Roshan Pillai, Keri Ramsey, Raj Gupta, Molly Brzezinski, Pooja Venugopal, Lorida Llaci, Brittany Gerald, Gabrielle Mills, Newell Belnap, Meredith Sanchez-Castillo, Chris D. Balak, Ana M. Claasen, Szabolcs Szelinger, Wayne M. Jepsen, Ashley L. Siniard, Ryan Richholt, Matt De Both, Marcus Naymik, Isabelle Schrauwen, Ignazio S. Piras, David W. Craig, Matthew J. Huentelman, Vinodh Narayanan, Sampathkumar Rangasamy

**Author notes:** Correspondence Dr. Sampathkumar Rangasamy, Neurogenomics Division, Translational Genomics Research Institute, Phoenix, Arizona, United States.

## Abstract

Mutations of the X-linked gene encoding methyl-CpG-binding protein 2 (*MECP2*) cause classical forms of Rett syndrome (RTT) in girls. Patients with features of classical Rett syndrome, but do not fulfill all the diagnostic criteria (e.g. absence of a *MECP2* mutation), are defined as atypical Rett syndrome. Genes encoding for cyclin-dependent kinase-like 5 (*CDKL5*) and forkhead box G1 (FOXG1) are more commonly found in patients with atypical Rett syndrome. Nevertheless, a subset of patients who are recognized to have an overlapping phenotype with RTT but are lacking a mutation in a gene that causes typical or atypical RTT are described as having Rett syndrome like phenotype (RTT-L). Whole Exome Sequencing (WES) of 8 RTT-L patients from our cohort revealed mutations in the genes *GABRG2, GRIN1, ATP1A2, KCNQ2, KCNB1, TCF4, SEMA6B*, and *GRIN2A*, which are seemingly unrelated to Rett syndrome genes. We hypothesized that the phenotypic overlap in RTT and RTT-L is caused by mutations in genes that affect common cellular pathways critical for normal brain development and function. We annotated the list of genes identified causing RTT-L from peer-reviewed articles and performed a protein-protein interaction (PPI) network analysis. We also investigated their interaction with RTT (typical or atypical) genes such as *MECP2, CDKL5, NTNG1*, and *FOXG1*. We found that the RTT-L-causing genes were enriched in the biological pathways such as circadian entrainment, the CREB pathway, and RET signaling, and neuronal processes like ion transport, synaptic transmission, and transcription. We conclude that genes that significantly interact with the PPI network established by RTT genes cause RTT-L, explaining the considerable feature overlap between genes that are indicated for RTT-L and RTT.

## Introduction

Rett syndrome (RTT), first described by Austrian neurologist Andreas Rett, is a deleterious X-linked neurodevelopmental disease mainly affecting females, which is characterized by the loss of spoken language and loss of purposeful hand use to hand stereotypies that mimic hand-washing^1^. Mutations of the X-linked gene encoding methyl-CpG-binding protein 2 (*MECP2*) causes RTT in girls, severe encephalopathy in male infants, and X-linked mental retardation^2^. Classical Rett Syndrome is mostly attributed to *de novo* mutations in *MECP2*^3^, with affected probands displaying a characteristic phenotype, including loss of acquired purposeful hand skills, loss of acquired spoken language, gait abnormalities, and stereotypic hand movements^4^. Patients who do not fulfill all the diagnostic criteria of RTT but presence of overlapping phenotype of RTT are classified as having atypical Rett syndrome^5^. Several pathogenic variants for atypical RTT have been described in recent years, such as the early-seizure onset variant attributed gene *CDKL5* (*cyclin-dependent kinase-like 5*)^6^, the preserved speech variant attributed to *MECP2*^7,8^, the rare variant attributed to gene *NTNG1* (*netrin-G1)*^9^, and the congenital variant attributed to gene *FOXG1* (*Forkhead box G1*)^10^.

Mutations also occur in loci other than *MECP2, CDKL5, NTNG1*, and *FOXG1*, with patients displaying some but not all the clinical features associated with classic and atypical RTT^12^. A proband with this phenotype is classified as Rett Syndrome like phenotype (RTT-L)^12^. The use of next generation sequencing (NGS) has expanded the locus heterogeneity in individuals diagnosed with RTT-L, however, the molecular pathways and processes causing the RTT-L phenotype are still unknown.^12^ The identification of pathogenic mutations in genes of RTT-L probands previously not associated to RTT or neurodevelopmental disorders indicates the need to understand the molecular mechanism so that effective therapies can be developed.

We hypothesized that several of the mutated genes causing the RTT-L phenotype encode closely interacting proteins with RTT and atypical RTT proteins; then, the analysis of their interactions might reveal a common biological pathway implicated in RTT-L. We report the use of a family-based exome sequencing approach in a cohort of 8 families with clinical features of RTT-L and the identification of *de novo* mutations in genes outside of the usually studied genes *MECP2, CDKL5, FOXG1*, and *NTNG1*. We then demonstrate the interactions that occur within the protein network between *MECP2, CDKL5, FOXG1*, and *NTNG1*. Finally, after assembling a list of *de novo* mutations in genes that cause RTT-L from published reports (including our own *de novo* mutations), we establish the interconnectedness of many genes from our list in relation to the protein network established by *MECP2, CDKL5, FOXG1*, and *NTNG1*.

## Materials and Methods

### Patient samples

In this study, we identified a cohort of 8 Caucasian trios with RTT-L clinical features according to the revised RettSearch International Consortium criteria and nomenclature,^4^ and all lacked mutations in *MECP2, CDKL5, FOXG1*, and *NTNG1*. The Rett Search International Consortium criteria is an instrument for assessing and diagnosing RTT. It classifies characteristics of RTT into typical and atypical RTT through a series of behavioral criteria. The parents of the affected children did not exhibit clinical features of RTT-L or intellectual disability. Genomic DNA from these trios was obtained from peripheral blood leukocytes. The study protocol and consent procedure were approved by the Western Institutional Review Board (WIRB; study number 20120789). Informed consent was obtained from patients.

### Whole exome sequencing

A trio-based WES method was performed. Genomic DNA was extracted from blood leukocytes for each member of the family trio and genomic libraries were prepared using 1.2 μg of DNA with the TruSeq DNA sample preparation and Exome Enrichment kit (62Mb; Illumina, San Diego, CA). Sequencing was performed by paired-end sequencing on a HiSeq2000 instrument (Illumina Inc, San Diego, USA) and were then aligned to the Human genome (Hg19/GRC37) using the Burrows-Wheeler alignment tool (BWA-MEMv0.7.8). Polymerase chain reaction (PCR) duplicates were removed using Picard (v1.128), and base quality recalibration and indel realignment were performed using the Genome Analysis Toolkit (GATKv3.3). Variants were called jointly with HaplotypeCaller^13^, recalibrated with GATK, and annotated with dbNSFP (v3.1) and snpEff (3.2a) for protein-coding events. Our WES was focused on *de novo* single nucleotide variants since they are often implicated in causing intellectual disability-related diseases, although other modes of inheritance such as autosomal recessive and X-linked were considered as well. To select variants that were not present in the healthy human populations, variants with an allele frequency >1% observed in the Single Nucleotide Polymorphism database (dbSNP) and the 1000 Genomes Pilot Project^14^ were filtered out and were filtered against variants found in their parents. Finally, *de novo* candidates were selected based on the alignment quality, damage predictors, and conservation level of each of the genes. The presence of the variants in large control databases such as the Genome Aggregation Database (gnomAD) was also evaluated^15^. *De novo* variants in subsequent analyses were defined as variants demonstrated by both exome sequencing and Sanger sequence validation to be present in a proband and absent from both parents. The predicted functional impact of each candidate’s *de novo* missense variant was assessed through analysis of *in silico* tools. Variants were considered potentially deleterious through varied criteria, including the MutationAssessor, MutationTaster, Provean, Combined Annotation Dependent Depletion (CADD), and Polyphen score, and splice-altering predictions for splice sites^16^.

### RTT-L Gene Literature Search

We manually curated a list of genes implicated in RTT-L. An extensive literature search was conducted on PubMed for peer-reviewed articles describing patients with RTT-L Syndrome or RTT Syndrome-like disorder till July 2017. We focused on genes where the literature clearly described the patient’s phenotypic features as overlapping with RTT Syndrome. The list was meant to be exhaustive, and as a result was routinely updated; the genes identified from our own study were included in the list. We also compiled a list of phenotypic characteristics observed in the subjects of the peer-reviewed articles we found. These characteristics were based off the criteria outlined by the RettSearch International Consortium as well as frequently described proband phenotypes^4^.

### Functional Enrichment Analysis and Ontology Lists

Functional enrichment analysis was performed on the RTT-L gene list along with RTT- and atypical RTT-causing genes (RTT genes) to infer significant biological processes and pathways. We used the Overrepresentation Test^17^ followed by the Fisher’s exact test (http://pantherdb.org, Version 13) based on the GeneOntology (GO) database^18^ (released 2017-12-27). Biological processes with the most significant overrepresentation were recorded, and the genes composing them were compiled into process-level gene lists. The Combined RTT-L and RTT gene lists’ involvement in biological pathways were analyzed using GeneAnalytics (http://geneanalytics.genecards.org) program to characterize molecular pathways, powered by GeneCards, LifeMap Discovery, MalaCards, and Path Cards^19^. Related pathways from the gene list were grouped into Superpathways to improve inferences and pathway enrichment, reduce redundancy, and rank genes within a biological mechanism via the GeneAnalytics algorithm. Pathways compiled were statistically overrepresented with significance defined at <0.0001 and a corresponding –log(p-value) score > 16.

### RTT- and RTT-L-Implicated Genes Network Analysis

Protein-protein interaction (PPI) networks were generated from the *Homo sapiens* interaction database created by GeneMania 2017-03-14 release^10^. The RTT-L-causing genes identified through our institution and our exhaustive literature search as well as the RTT-implicated genes (*MECP2, CDKL5, FOXG1, NTNG1*) were used as the seed list of a PPI network of 250 total genes. Network figures were created using Cytoscape 3.5.1^21^. PPI network statistics were generated using Cytoscape’s NetworkAnalyzer^22^ and Centiscape^23^. To estimate the statistical significance of observations drawn from gene list PPI networks, we compiled a gene list of 1,460 highly expressed central nervous system (CNS) genes identified through “The Human Protein Atlas”^24^ (www.proteinatlas.org). For each biological process and pathway gene list identified, PPI networks containing the 250 genes, the statistically significant threshold were created with a seed of random brain genes (n=65) from The Human Protein Atlas gene list. The mean numbers and standard deviation of network centralization, network density, node degree, and edge length of the random gene lists were used to calculate z-scores of RTT-L and RTT statistics (Figure 1).

**Figure 1:**
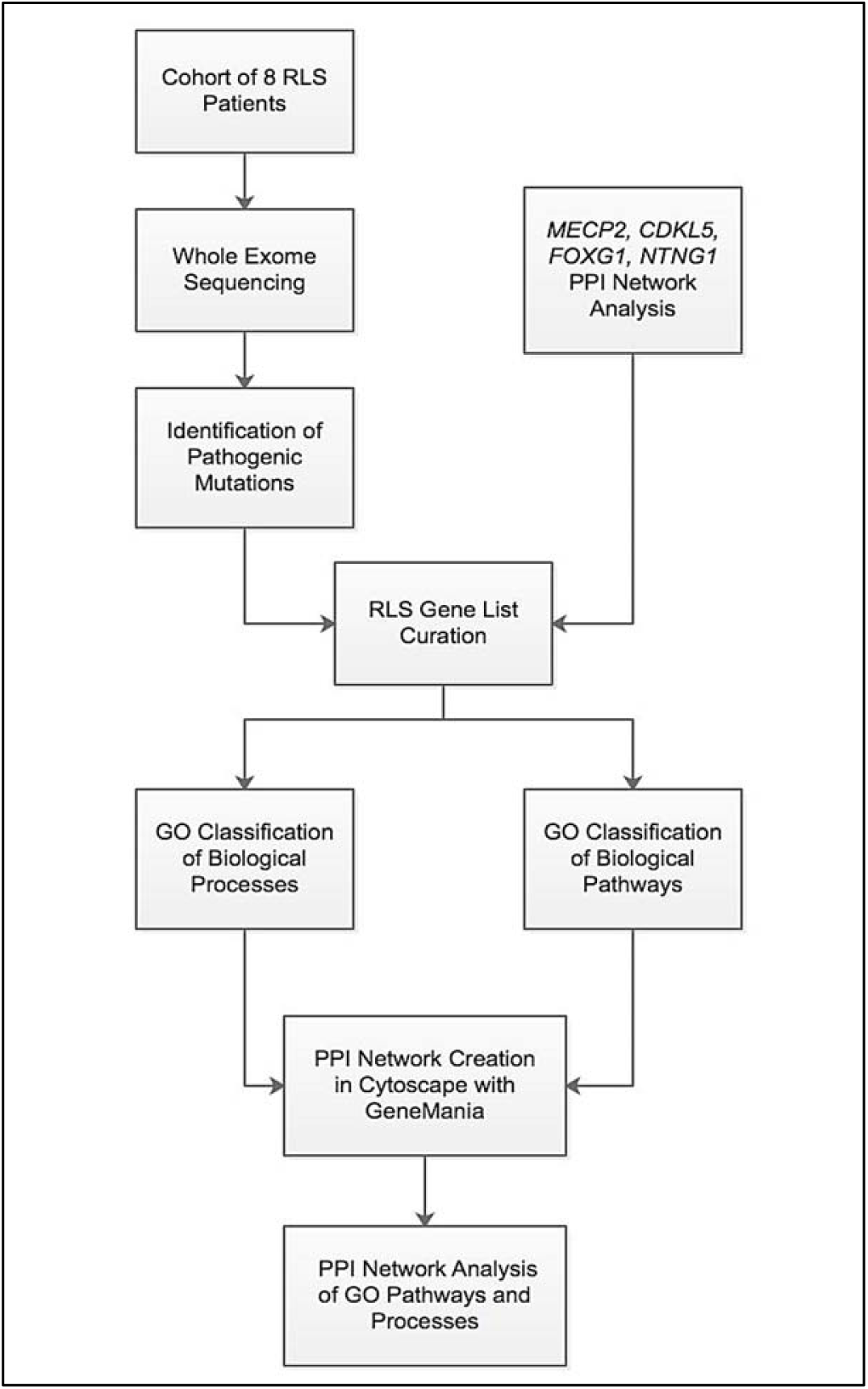
Flow diagram of the data analysis.

## Results

### Clinical and Molecular Characterization of Pathogenic Variants

Here we present the summary of the clinical reports behind each RTT-L case at our institution (Table 1). All these patients were initially diagnosed as RTT or RTT-L by the clinician before the genetic diagnosis. Through WES, we identified disease causing variants in *GABRG2* (patient 1), *GRIN1* (patient 2), *ATP1A2* (patient 3), *KCNQ2* (patient 4), *KCNB1* (patient 5), *GRIN2A* (patient 6), *TCF4* (patient 7), and *SEMA6B* (patient 8) genes, all of which occurring in evolutionarily conserved locations (Figure 2). In addition, all variants were not observed in controls in the Genome Aggregation Database (gnomAD). These genes presented phenotypes that overlap with those seen in patients with Rett syndrome (Table 1). Patient 1 is heterozygous for a *de novo* substitution mutation (c.316G>A) in the *GABRG2* gene, which is associated with childhood febrile seizures, resulting in a missense (p.Ala106Thr) in the ligand binding region. Patient 2 was identified to carry a heterozygous *de novo* mutation in *GRIN1* gene, which is associated with neurodevelopmental disorder with seizures. The heterozygous c.2443G>C substitution resulted in a missense mutation (p.Gly815Arg). Patient 3 is heterozygous for a *de novo* substitution mutation (c.977T>G) (p.Ile326Arg) in the *ATP1A2* gene, which is associated with childhood hemiplegia. Trio WES performed on Patient 4 revealed a *de novo* substitution mutation (c.740C>A) in the gene *KCNQ2*, which is associated with early infantile epileptic encephalopathy, resulting in a nonsense variant (p.Ser247Ter), which is predicted to be targeted by nonsense-mediated decay. Patient 5 was identified to have a *de novo* variant (c.916C>T) in *KCNB1 gene*, which is associated with early infantile epileptic encephalopathy, resulting in a missense (p.Arg306Cys). Patient 6 is heterozygous for a *de novo* mutation (c.1845T>C) in gene *GRIN2A*, which encodes a member of the glutamate-gated ion channel protein family and is associated with focal epilepsy and mental retardation. This mutation results in missense (p.Asn614Ser) in exon 8 and affects the ligand-gated ion channel domain in the cytoplasmic region. Patient 7 is heterozygous for a *de novo* splice site mutation in *TCF4* gene. WES of Patient 8 revealed the presence of a *de novo* frameshift (c.1991delG) in the gene *SEMA6B*, which is predicted to led to premature truncated protein (p.Gly664fs).

**Table 1:** Clinical summary of and molecular characterization of patients in our RTT-L cohort.

| Patient | Developmental Regression | Developmental Delay | Intellectual Disability | Microcephaly | Loss of Hand Use | Stereotyped Hand Movements | Involuntary Tongue Movements | Hyperventilation | Choreoathetosis | Early Epileptic Encephalopathy | Hypotonia | Scoliosis | Gene | Protein | Variant: genomic coordinates | cDNA change | Protein change | Gene-diseases association |
| --- | --- | --- | --- | --- | --- | --- | --- | --- | --- | --- | --- | --- | --- | --- | --- | --- | --- | --- |
| 1 | Y | Y | Y | Y | N | N | N | N | N | Y | Y | N | GABRG2 | Gamma-Aminobutyric Acid Type A Receptor Gamma2 Subunit | 5:161522557 | c.316 G>A | p.Ala106Thr | Epilepsy, generalized, with febrile seizures plus, type 3 |
| 2 | N | Y | Y | Y | N | N | N | Y | Y | N | Y | N | GRIN1 | Glutamate Ionotropic Receptor NMDA Type Subunit 1 | 9:140058120 | c.2443 G>C | p.Gly815Arg | Mental retardation, autosomal dominant 8 |
| 3 | Y | Y | Y | Y | Y | Y | N | N | Y | Y | Y | N | ATP1A2 | ATPase Na<sup>+</sup>/K<sup>+</sup> Transporting Subunit Alpha 2 | 1:160097570 | c.977 T>G | p.Ile326Arg | Alternating hemiplegia of childhood |
| 4 | N | Y | Y | N | N | N | N | N | Y | Y | N | N | KCNQ2 | Potassium Voltage-Gated Channel Subfamily Q Member 2 | 20:63442482 | c.740 C>A | p.Ser247Ter | Epileptic encephalopathy, early infantile, 7 |
| 5 | Y | Y | Y | N | N | N | N | N | N | N | N | N | KCNB1 | Potassium Voltage-Gated Channel Subfamily B Member 1 | 20:47991181 | c.916 C>T | p.Arg306Cys | Epileptic encephalopathy, early infantile, 26 |
| 6 | N | Y | N | N | N | Y | N | N | Y | N | N | Y | GRIN2A | Glutamate Ionotropic Receptor NMDA Type Subunit 2A | 16:9923446 | c.1845 T>C | p.Asn614Ser | Epilepsy, focal, with speech disorder and with or without mental retardation |
| 7 | Y | Y | Y | Y | Y | Y | Y | Y | N | N | Y | N | TCF4 | Transcription Factor 4 | chr18:52901774 | c.1486+5 G>T | IVS16+5 G>T (Splice Variant) | Pitt-Hopkins |
| 8 | N | Y | Y | Y | N | Y | N | N | N | N | Y | N | SEMA6B | Semaphorin 6B | chr19:4544288 | c.1991delG | p.G664AfsX21 | Autism Spectrum Disorder |

**Figure 2:**
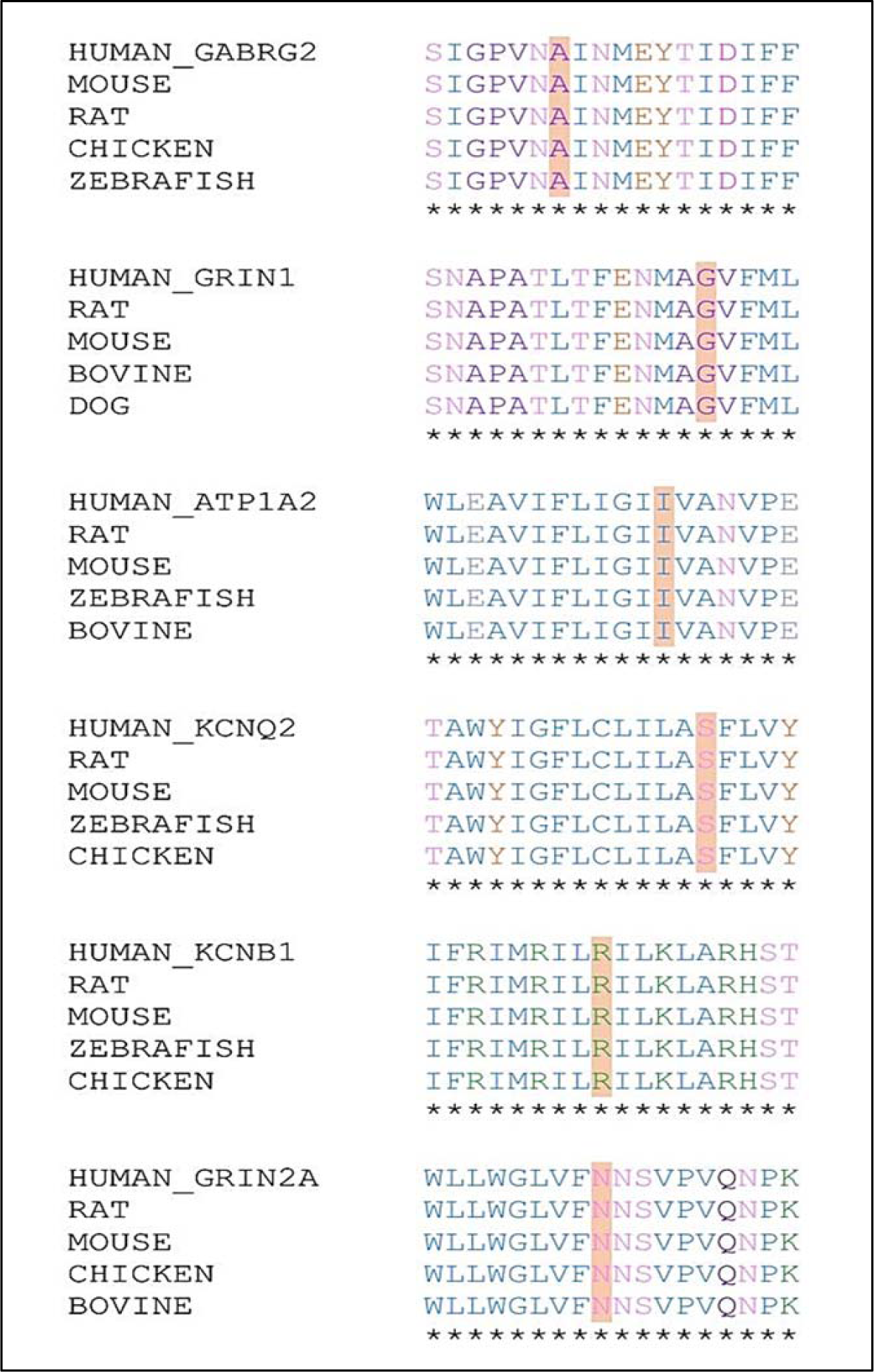
UniProt alignment of the RTT-L genes identified by our cohort. Mutation location sites are identified by orange highlight.

### Curation of Gene Lists

After observing phenotypic overlap between features of RTT and our patients, we curated a list of genes implicated in causing RTT-L from articles in peer-reviewed journals^12,25-40^. Through an exhaustive literatures search, 58 genes were identified to have *de novo* damaging or chromosomal deletion mutations that caused RTT-L (Table 2). In addition to the curated list, we included the genes (*GABRG2, GRIN1, ATP1A2, KCNB*, and *GRIN2A*) identified from our cohort plus RTT-causing genes such *MECP2, CDKL5, FOXG1*, and *NTNG1* (Complete RTT-L and RTT Gene List).

**Table 2:**
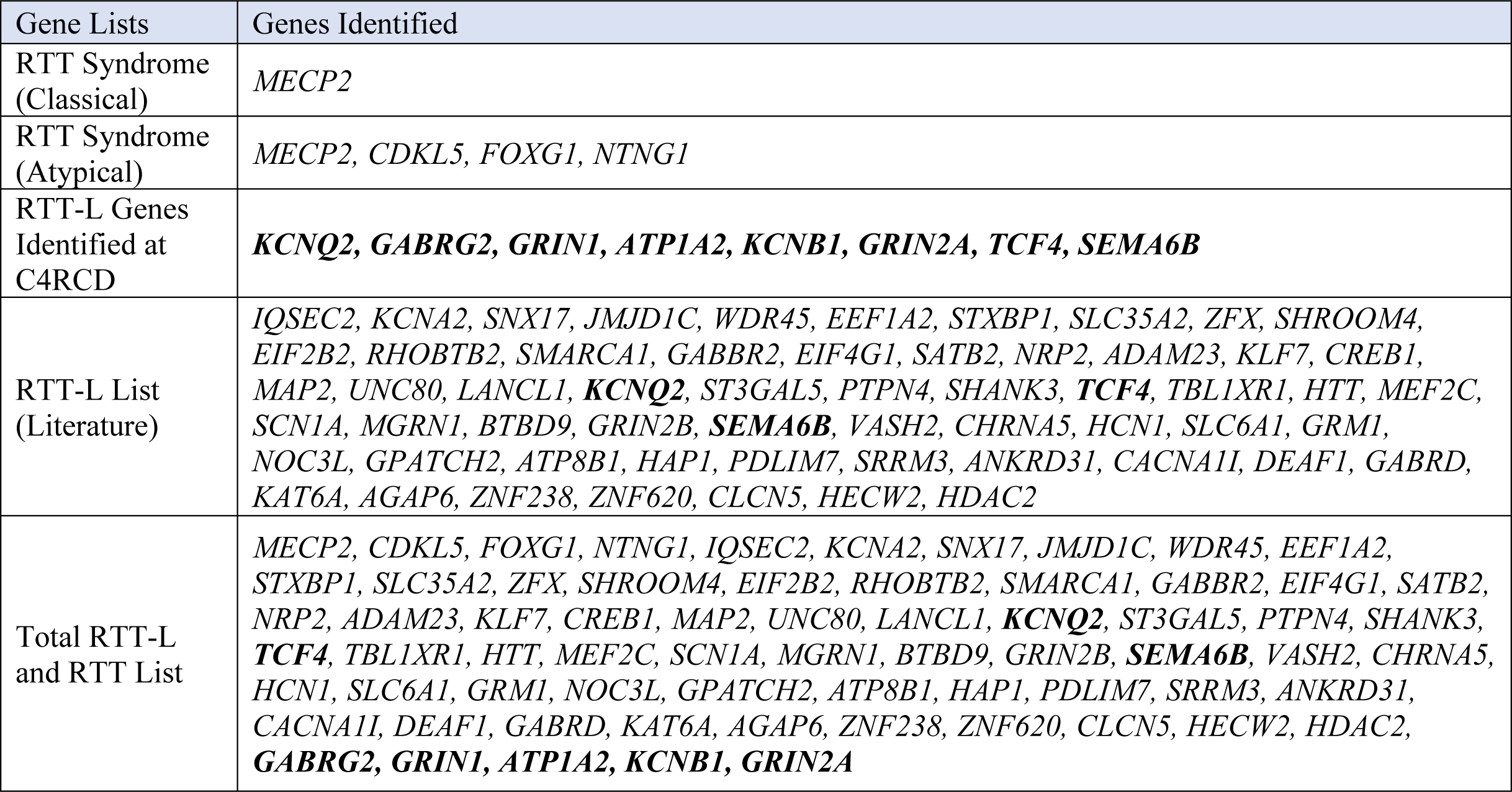
List of genes contributing to classical RTT, atypical RTT, RTT-L features of our cohort, RTT-L, and both RTT-L and RTT. Bolded genes indicate those identified in our patient cohort as harboring pathogenic mutations.

### Phenotype Clustering of RTT-L Patients

We analyzed the phenotypes of RTT-L patients to evaluate the frequency of main and supportive RTT criteria that appeared in patients. Of the main criteria, RTT-L patients often displayed at least two of the following clinical RTT phenotype: regression, partial or complete loss of acquired purposeful hand skills, partial or complete loss of acquired spoken language, gait abnormalities, and hand stereotypies. However, RTT-L patients often frequently displayed Rett supportive criteria, including bruxism (56%), growth retardation (67%), respiratory disturbances (43%), and screaming fits (33%).

### Functional Enrichment

We then conducted functional enrichment using GeneAnalytics on the complete RTT-L and RTT gene list to identify the overrepresented or under-represented biological processes and pathways. The biological functions identified (p < 0.0001) were ranked according to the p-value (Table 3). RTT-L patients showed significant over-representation of GO Biological Processes involving ion transport, synaptic transmission, regulation of ion transport, visual learning, regulation of postsynaptic membrane potential (Table 3). Similarly, RTT-L patients also had over-representation of the GeneAnalytics Pathways including Synaptic transmission, dopamine signaling, neurophysiological Process, NMDA receptor activation and circadian entrainment (Table 3). The identification of these biological pathways and processes suggests the diversity of biological pathways and processes RTT-L genes are implicated in.

**Table 3:**
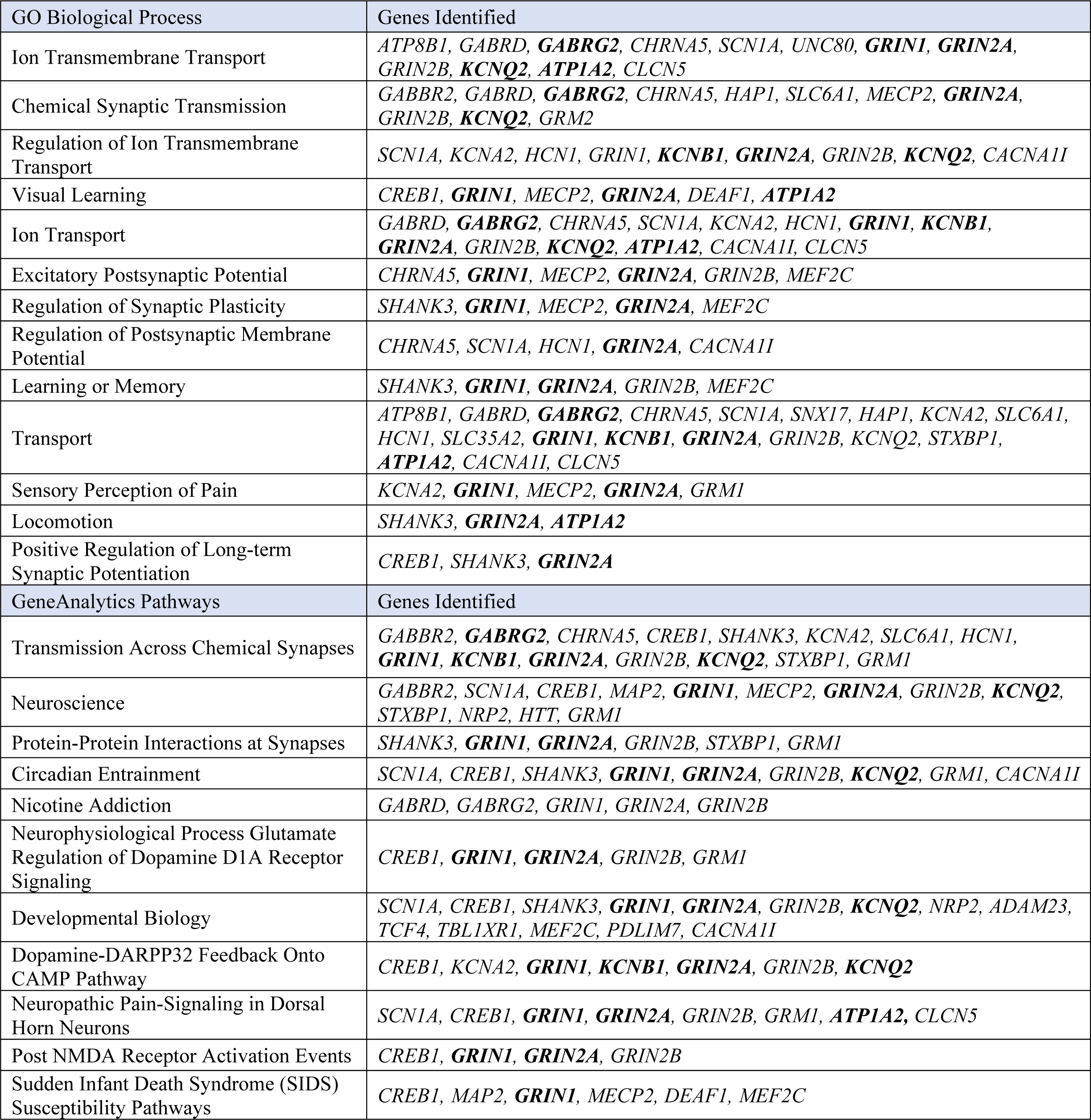
Lists of genes associated with each GeneOntology biological process and GeneAnalytics Pathways. Bolded genes indicate those identified in our patient cohort as harboring pathogenic mutations.

### Comparison of RTT-L-Causing Genes with SFARI Autism Gene Set

In 2009, Wall et al. created a phylogeny^41^ which grouped autism with RTT Syndrome under the term “autism sibling disorders”. The authors used the SFARI gene database to collect 991 genes implicated in autism. In order to evaluate our data, we compared the genes implicated in causing RTT-L with the SFARI autism gene set. The SFARI gene set showed a significant number of genes in common with the curated gene list. 37% of the genes in the RTT-L and RTT were implicated in the autism-causing SFARI gene set (Supplementary Table 1).

### Network Analysis of Protein-Protein Interactions Between RTT-L and RTT Genes

Phenotypic overlap between clinical features of RTT and the RTT-L patients in our cohorts was observed, prompting an investigation of protein-protein interactions of genes implicated in RTT-L Syndrome (Table 1). We used the GeneMania interaction data sets to examine whether *de novo* mutations causing RTT syndrome and RTT-L syndrome clustered in protein-protein interaction networks. Remarkably, 63 of the 67 genes mapped to an interconnected PPI network with 351 edges, with only the genes *SHANK3* and *ZNF620* being disconnected from the network. These 63 genes interacted with one another through coexpression (45.47%), predicted interactions (17.20%), pathway interactions (12.34%), colocalization (11.72%), physical interactions (9.61%), shared protein domains (2.52%), and genetic interactions (1.15%) (Figure 3A). This PPI network was then supplemented with the addition of 183 genes, determined as having the strongest interaction with the 67 RTT and RTT-L genes curated through the GeneMania association algorithm; the modified PPI network (n=250) was analyzed for network centralization, average node degree, average edge length, and network density (Table 4). The PPI network had significantly high average node degree (p=0.0003), network density (p=0.0007), and network centrality (p=0.0015) compared to control networks of brain genes identified through Human Protein Atlas, contributing to the greater connectivity of the network in shared neighbors between nodes (Table 4 and Figure 3B). The network was also characterized by the presence of genes (*FOXG1, CAMK2D, GFRA1, DAB1, MEF2C, PIK3R1, PTPRT, SNAP25, MAP3K10, SH3GL2, SYT1*) with high betweenness centrality, indicating that these particular genes play an outsized role in connecting different biological pathways and processes together (Figures 3C and 3D).

**Table 4:** Network analysis of PPI networks generated from the seed gene list.

| Gene List | Number of Seed Nodes | Centralization | Avg. Node Degree | Avg. Edge Length | Density |
| --- | --- | --- | --- | --- | --- |
| RTT-L List (Literature) | 61 | 0.293 | 51.744 | 1.84 | 0.208 |
| Complete RTT-L and RTT List | 65 | 0.274 | 54.312 | 1.824 | 0.218 |
| GO Biological Process | Number of Seed Nodes | Centralization | Avg. Node Degree | Avg. Edge Length | Density |
| Ion Transmembrane Transport | 12 | 0.228 | 63.672 | 1.765 | 0.256 |
| Chemical Synaptic Transmission | 11 | 0.285 | 63.56 | 1.764 | 0.255 |
| Regulation of Ion Transmembrane Transport | 9 | 0.274 | 54.312 | 1.824 | 0.218 |
| Visual Learning | 6 | 0.284 | 12.952 | 2.726 | 0.052 |
| Ion Transport | 14 | 0.211 | 64.864 | 1.754 | 0.26 |
| Excitatory Postsynaptic Potential | 6 | 0.286 | 40.264 | 1.951 | 0.162 |
| Regulation of Synaptic Plasticity | 5 | 0.336 | 35.104 | 1.926 | 0.141 |
| Regulation of Postsynaptic Membrane Potential | 5 | 0.409 | 38.936 | 1.908 | 0.156 |
| Learning or Memory | 5 | 0.486 | 38.912 | 1.887 | 0.156 |
| Transport | 20 | 0.258 | 53.304 | 1.827 | 0.214 |
| Sensory Perception of Pain | 5 | 0.376 | 41.168 | 1.902 | 0.165 |
| Locomotion | 3 | 0.488 | 38.416 | 1.874 | 0.154 |
| Positive Regulation of Long-term Synaptic Potentiation | 3 | 0.608 | 28.76 | 1.946 | 0.116 |
| GeneAnalytics Pathways | Number of Seed Nodes | Centralization | Avg. Node Degree | Avg. Edge Length | Density |
| Transmission Across Chemical Synapses | 15 | 0.246 | 56.16 | 1.806 | 0.226 |
| Neuroscience | 13 | 0.248 | 29.768 | 2.182 | 0.12 |
| Protein-Protein Interactions at Synapses | 6 | 0.363 | 27.296 | 2.205 | 0.11 |
| Circadian Entrainment | 9 | 0.271 | 51.048 | 1.868 | 0.205 |
| Nicotine Addiction | 5 | 0.579 | 55.944 | 1.789 | 0.225 |
| Neurophysiological Process Glutamate Regulation of Dopamine D1A Receptor Signaling | 5 | 0.419 | 34.504 | 1.933 | 0.139 |
| Developmental Biology | 14 | 0.294 | 49.488 | 1.849 | 0.199 |
| Dopamine-DARPP32 Feedback Onto CAMP Pathway | 7 | 0.243 | 48.976 | 1.862 | 0.197 |
| Neuropathic Pain-Signaling in Dorsal Horn Neurons | 8 | 0.266 | 41.312 | 1.921 | 0.166 |
| Post NMDA Receptor Activation Events | 4 | 0.393 | 31.904 | 1.944 | 0.128 |
| Sudden Infant Death Syndrome (SIDS) Susceptibility Pathways | 6 | 0.287 | 21.072 | 2.193 | 0.085 |

**Figure 3:**
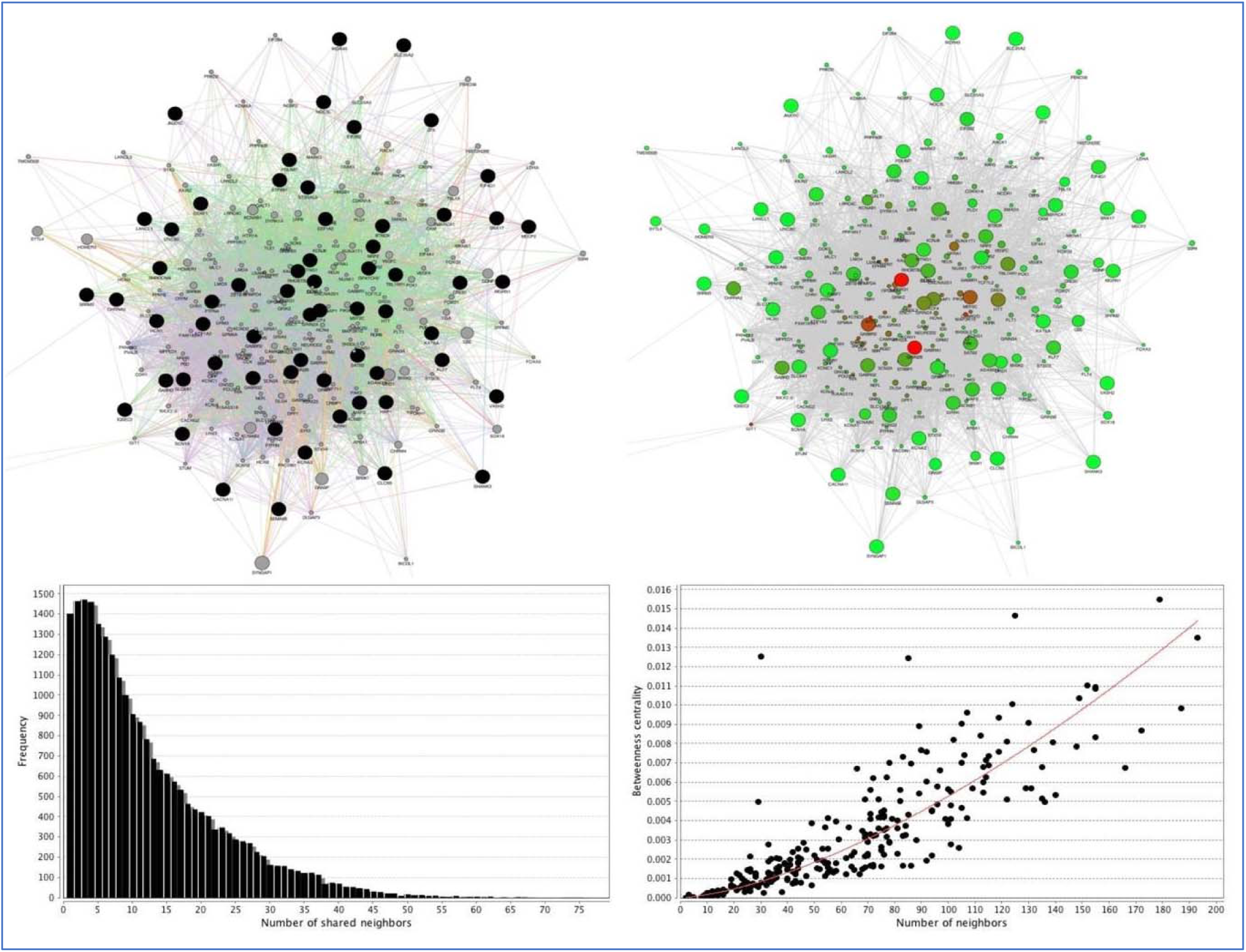
(A) A PPI network around the RTT-L genes. Nodes in the network correspond to genes, and edges correspond to interactions. Node sizes are proportional to the gene’s contribution to the overall cluster score. Edge widths are proportional to the strength of interaction between the genes. Edges are colored based on the type of interaction between genes: green indicates genetic interactions, red indicates physical interactions, purple indicates coexpression, dark blue indicates colocalization, yellow indicates shared protein domains, orange indicates predicted interaction, and light blue indicates shared pathway interactions. (B) The PPI network shown in (A) with nodes colored by degree of betweenness centrality. (C) Frequency of nodes that have a given number of shared neighbors. (D) A plot of betweenness centrality of the nodes in the network as a function of their number of neighbors. The close correlation of an exponential line of best fit shows the presence of genes that harbor interactions with many other different genes that serve as the focal point of interactions between different biological processes and pathways.

### RTT-L Gene Interaction with MECP2 Epigenetic Network

Most frequently associated with causing classical Rett syndrome, *MECP2* forms a complex regulatory network, regulating neuronal function and development by regulating RNA splicing and chromatin structure as well as interfering in methylation of DNA. We analyzed the *MECP2* regulatory network from Ehrhart et. al., for genes implicated in RTT-L genes and pathways. We found that RTT-L genes were often directly involved in the *MECP2* regulatory network. *MECP2* activates the formation of the MECP2-HDAC complex, which in turn inhibits *MEF2C*, another RTT-L gene. *MECP2* also forms complexes with RTT-L gene *CREB1* to activate transcription as well as with *YB1* and *PRPF3* to regulate the alternative splicing of RTT-L gene *GRIN1*. Additionally, *MECP2* affects the glutamate and GABA pathways, both of which contain many RTT-L genes, through the activation of *AMPA* and the inhibition of noncoding RNA Ak081227, respectively.

## Discussion

Recent publications described the application of WES to patients with features of RTT, identifying several variants beyond those found in *MECP2, CDKL5, FOXG1*, and *NTNG1*. As a result, significant questions like which genes could cause a RTT-L phenotype, mutations in which biological pathways and processes lead to clinical RTT-L features, and what is the structure of the PPI networks associated with the RTT-L phenotype remain unanswered. We attempted to answer these questions through: i) extensive analysis of published literature and from WES on a group of eight trios from our center whose probands displayed RTT-L features, the functional enrichment of genes implicated in RTT-L, and ii) the PPI network analysis of gene lists distinguished by overrepresented biological functions and processes.

WES was performed on the eight trios in our RTT-L cohort, and eight unique *de novo* variants in genes that had no previous association with classical or atypical RTT: *GABRG2, GRIN1, GRIN2A, KCNB1, KCNQ2, ATP1A2, TCF4*, and *SEMA6B*. The phenotypes associated with these genes overlap with those observed in our cohort of RTT-L patients. Combined analysis of the RTT-L cohort indicates a mutational heterogeneous profile hitting shared neurological processes and pathways contributes to RTT-L features, especially when directly interacting with the RTT-implicated genes *MECP2, CDKL5, FOXG1*, and *NTNG1*. Examples of functional overlap are potassium voltage-gated channel subfamily B member 1 (*KCNB1*), which is responsible for transmembrane potassium transport in excitable membranes, and potassium voltage-gated channel subfamily Q member 2 (*KCNQ2*), responsible for the subthreshold electrical excitability of neurons as well as the responsiveness to synaptic inputs. These transmembrane ion proteins are functionally relevant in the context of RTT and RTT-L, with *KCNB1* implicated in earlier studies of RTT-L. Other examples of functional overlap are glutamate ionotropic receptor NMDA type subunit 1 (*GRIN1*) and glutamate ionotropic receptor NMDA type subunit 2A (*GRIN2A*), both of which are essential in excitatory neurotransmission, verbal memory, and cognitive function. Additionally, the identification of gamma-aminobutyric acid type A receptor gamma2 subunit (*GABRG2*) contributes to the characterization of GABA receptors as implicated in causing RTT-L features. Finally, the implication of ATPase Na+/K+ transporting subunit alpha 2 (*ATP1A2*) indicates the links between hemiplegia and RTT, particularly with regard to control of voluntary motion. These findings indicate the significant degree of overlap between RTT-L and other neurodevelopmental disorders. Our WES indicated that patients displaying RTT-L features carried variants in well-known genes known to cause childhood epilepsy (*KCNB1, KCNQ2, GABRG)*, mental retardation (*GRIN1, GRIN2A*), and hemiplegia (*ATP1A2*).

The creation of a list of genes implicated in RTT-L combined with functional enrichment further demonstrates the mutational diversity of the clinical RTT phenotype. Past studies have noted genes causing RTT-L that have been implicated in neurodevelopmental diseases like Dravet syndrome (*SCN1A*), Pitt-Hopkins syndrome (*TCF4*), and Huntington’s disease (*HTT*). This clinical overlap with similar neurodevelopmental diseases further suggests the presence of shared functional networks. The performance of functional enrichment with GeneAnalytics and the Panther overrepresentation test further supports this idea. The identification of GO biological processes like ion transmembrane transport, chemical synaptic transmission, and regulation of ion transmembrane transport substantiated the notion that genes implicated in neuronal activity played a significant role in contributing to RTT-L features; however, the overrepresentation of biological processes of locomotion and memory emphasized the broad diversity of biological pathways RTT-L genes are implicated in (Supplementary Figure 1 & 2). Similarly, GeneAnalytics analysis detected somewhat expected overrepresentation of pathways like transmission across chemical synapses and protein-protein interactions at synapses as well as unprecedented pathways like circadian entrainment and nicotine addiction. The overrepresentation of these biological processes and pathways accentuates the broad range of pathways that contain RTT-L-implicated genes.

Network analysis of the PPI interaction network generated with the RTT and RTT-L syndrome-causing genes shows the presence of a significantly interconnected protein network facilitated by genes with high betweenness centrality, or the number of shortest paths in the network that pass through the gene. These genes serve as key interactors with genes that cause RTT-L syndrome. For instance, one of these high betweenness centrality genes, synaptotagmin 1 (*SYT1*), has protein-protein interactions with 20 of the 56 genes that cause RTT-L syndrome as well as with *CDKL5*. Moreover, the PPI network generated by these high-centrality genes contains many of the GO biological processes also implicated in the RTT-L PPI network.

Several of these overrepresented biological processes and pathways are critically involved in protein interactions involving classic and atypical RTT-causing genes. As previously mentioned, RTT-L-causing genes are directly involved in the *MECP2*-involved pathways of chromatin regulation (*HDAC2, MEF2C, CREB1, GRIN1*) and upregulation of glutamate and downregulation of dopamine and GABA pathways (*CREB1, KCNA2, GRIN1, KCNB1, GRIN2A, GRIN2B*, and *KCNQ2*)^42^. And while the RTT-L-causing genes do not play a role in the transcriptome profiling described by Lin et al., they share overlapping biological processes in many neuronal functions that have altered expression in RTT, such as synaptic transmission, neurotransmission, excitation of neurons, and development of neurons^43,44^. The involvement of many RTT-L-implicated genes in *MECP2* function indicates that the replication of many of clinical features of RTT-L similar to that of classical or atypical RTT could be attributed to the disruption of one of *MECP2*’s transcription regulatory activities. This interaction between *MECP2* and RTT-L implicated genes reinforces the understanding that there is not a unique biological process or molecular pathway involved in RTT that is disrupted; rather, because *MECP2* is involved in a variety of pathways by controlling transcription, as seen from Erhart et al., *MECP2* functionality can be interrupted by alterations in the many different pathways it controls.

The various diseases implicated in RTT-L syndrome-causing genes and diversity of clinical phenotypes complicates defining a set of features indicative of RTT-L Syndrome (Supplementary Figure 3). The phenotypes of the genes previously associated with NDDs and ASD but now implicated in RTT-L syndrome often overlap with those observed in our cohort of RTT-L patients. Based on the characteristics of our patient cohort and the diseases RTT-L syndrome genes are implicated in, most RTT-L patients tend to have clinical features of, from most to least frequent, epilepsy, intellectual disorder, regression, microcephaly, and hand stereotypies (Table 1). While the results of our phenotype clustering demonstrate many different combinations of these clinical features, the future clustering of these features could aid in the delineation of a central phenotype of RTT-L syndrome, as well as guide diagnoses for genetic testing in future patients.

In this study, we identified unreported variants in five novel genes that cause RTT-L syndrome. Further genetic characterizations of RTT-L patients are critical for future genotype-phenotype correlations and the establishment of a set of core RTT-L clinical features. Finally, our characterization of the biological processes and pathways implicated in RTT-L syndrome-causing genes showed the common pathways these genes converge to and offers guidance for the development of future target therapies. Finally, the analysis of PPI networks of RTT-L syndrome-causing genes identifies novel genes of high centrality that could be future candidate genes or be affected by genes that cause RTT-L syndrome.

This material is original research and has not been previously published. It has not been submitted for publication elsewhere while under consideration. The authors declare that there is no conflict of interest associated with this manuscript.

## Acknowledgements

The authors thank the family for participating in this study and all the previous members of the C4RCD research group not included in the author list.

## Consent

The study was explained to the extent compatible with the subject’s understanding and was enrolled into the Center for Rare Childhood Disorders program at the Translational Genomics Research Institute (TGen). The study protocol and consent procedure were approved by the Western Institutional Review Board (WIRB study number#: 20120789).

## Supplementary Materials

**Supplementary Figure 1:**
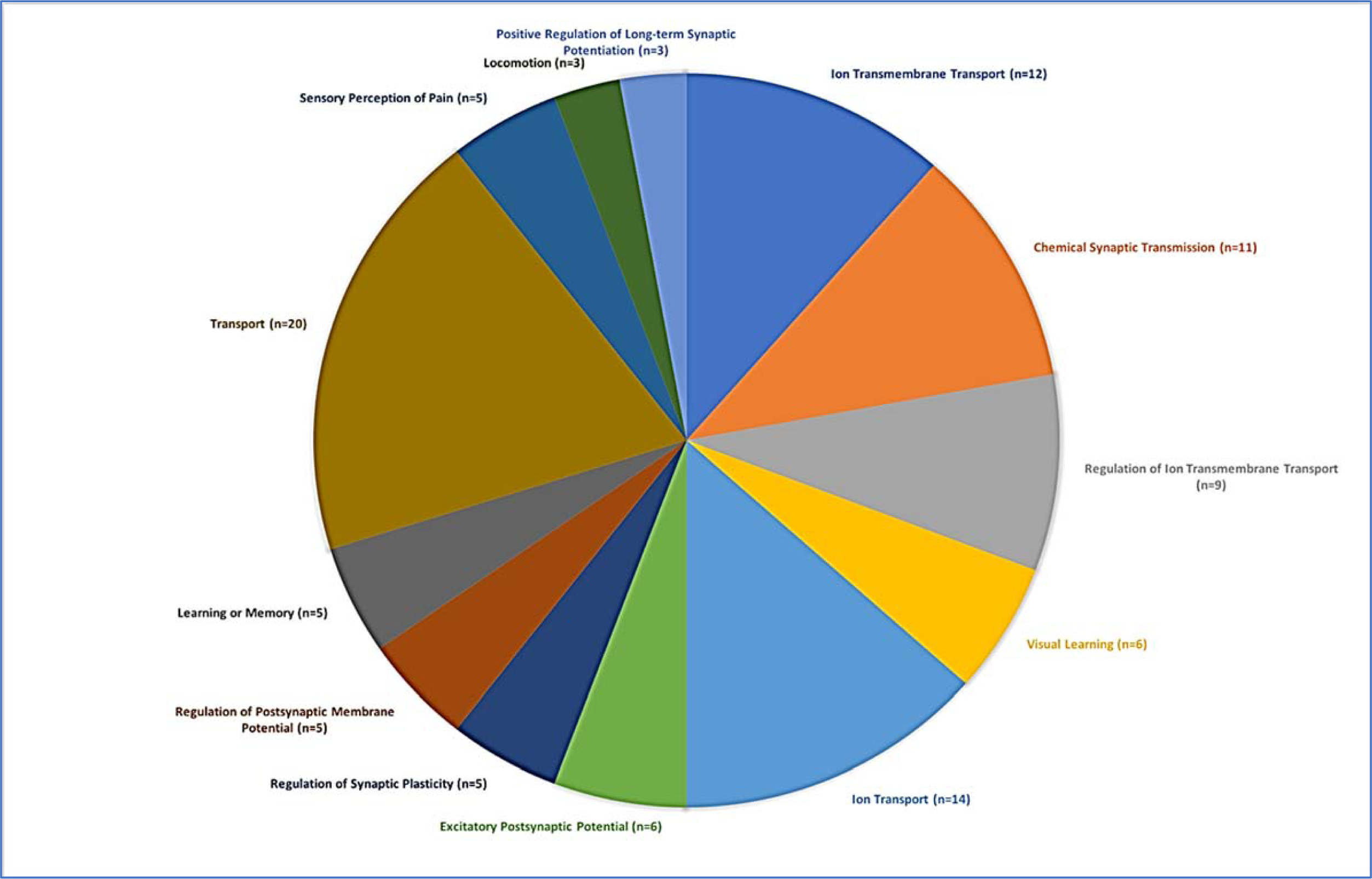
Number of RTT-L-causing genes associated with biological processes described by GeneOntology

**Supplementary Figure 2:**
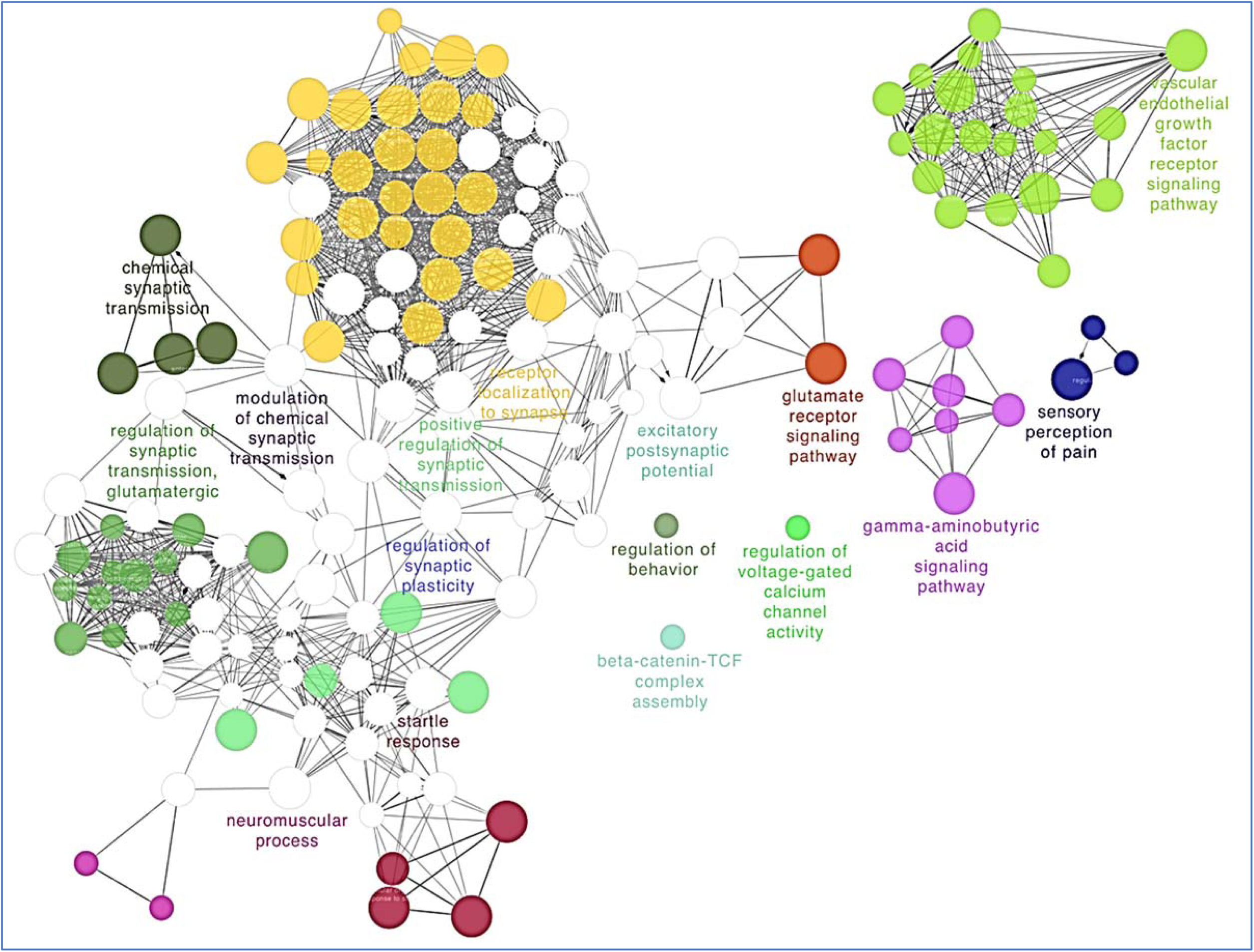
Pathway clustering of the varied biological processes of RTT-L-causing genes.

**Supplementary Figure 3:**
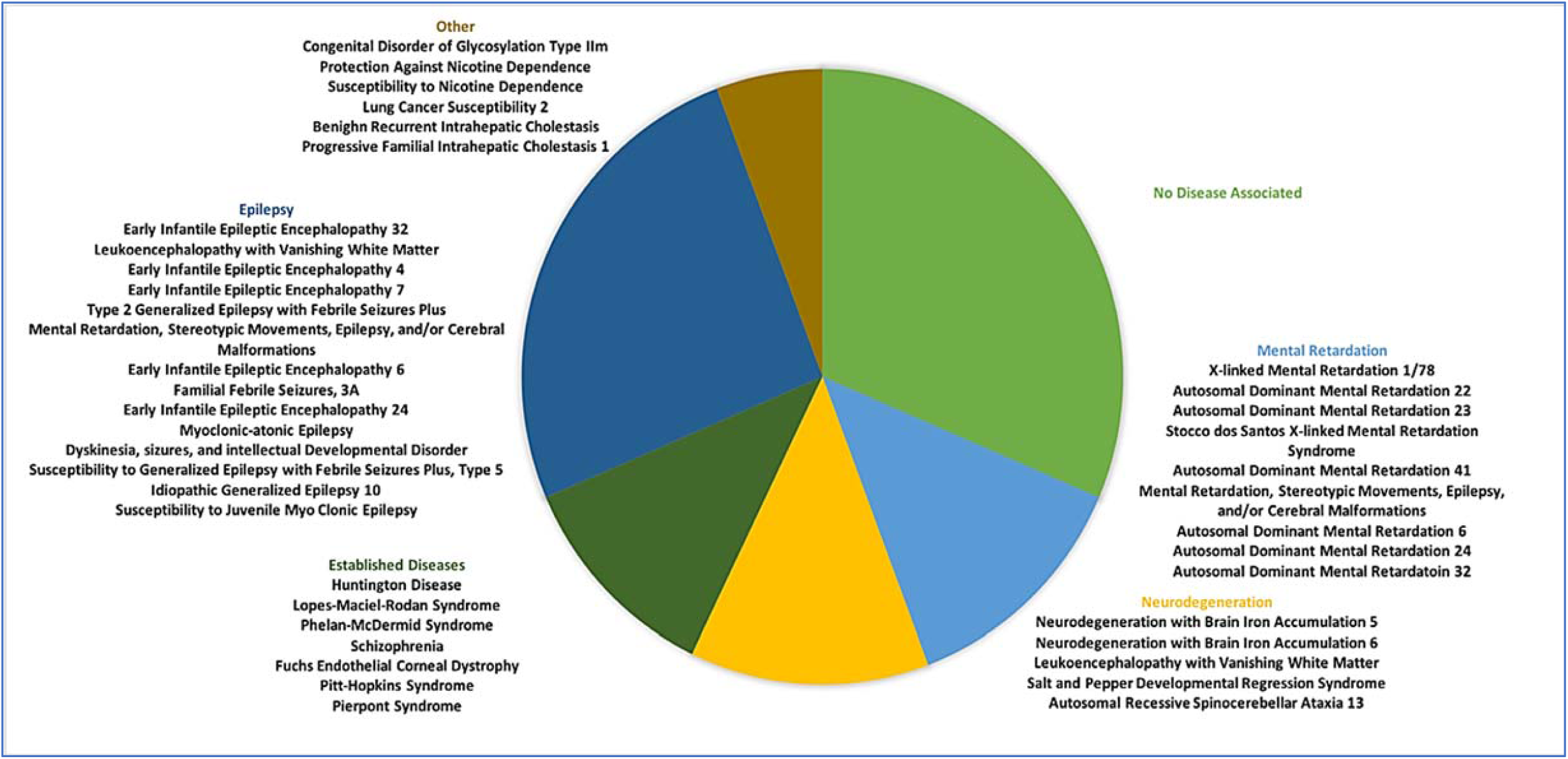
Online Mendelian Inheritance in Man (OMIM) diseases/disorders associated with different RTT-L-causing genes, grouped by disease class and function.

**Supplementary Table 1:** List of RTT-L-causing genes that are implicated in causing autism according to the Simons Foundation Autism Research Initiative (SFARI)

| **Autism**<br>**Genes** |
| --- |
| CACNA1I |
| DEAF1 |
| EEF1A2 |
| GRIN1 |
| GRIN2A |
| GRIN2B |
| GRM1 |
| HCN1 |
| IQSEC2 |
| JMJD1C |
| KAT6A |
| KCNB1 |
| KCNQ2 |
| MAP2 |
| MEF2C |
| NRP2 |
| SATB2 |
| SCN1A |
| SHANK3 |
| SLC6A1 |
| STXBP1 |
| TBL1XR1 |
| TCF4 |
| UNC80 |

